# MS-EmpiRe utilizes peptide-level noise distributions for ultra sensitive detection of differentially abundant proteins

**DOI:** 10.1101/514000

**Authors:** Constantin Ammar, Markus Gruber, Gergely Csaba, Ralf Zimmer

## Abstract

Mass spectrometry based proteomics is the method of choice for quantifying genome-wide differential changes of proteins in a wide range of biological and biomedical applications. Protein changes need to be reliably derived from a large number of measured peptide intensities and their corresponding fold changes. These fold changes vary considerably for a given protein.

Numerous instrumental setups aim to reduce this variability, while current computational methods only implicitly account for this problem. We introduce a new method, MS-EmpiRe (github.com/zimmerlab/MS-EmpiRe), which explicitly accounts for the noise underlying peptide fold changes. We derive dataset-specific, intensity-dependent empirical error distributions, which are used for individual weighing of peptide fold changes to detect differentially abundant proteins. The method requires only peptide intensities mapped to proteins and, thus, can be applied to any common quantitative proteomics setup. In a recently published proteome-wide benchmarking dataset, MS-EmpiRe doubles the number of correctly identified changing proteins at a correctly estimated FDR cutoff in comparison to state-of-the-art tools. We confirm the superior performance of MS-EmpiRe on simulated data. MS-EmpiRe provides rapid processing (< 2min) and is an easy to use, general-purpose tool.

## 2 Introduction

A major fraction of current Mass Spectrometry (MS) based proteomics experiments is quantitative in nature and aims at the detection and quantification of differentially expressed proteins between biological conditions [1]. As MS measurements are subject to substantial noise, researchers have to rely on statistical tests which detect changing proteins at a given false discovery rate (FDR). The de-facto value of a quantitative proteomics experiment could hence be defined by the overall sensitivity (i.e. the fraction of all changing proteins, which is actually detected by a statistical test) at a reasonable FDR. Huge instrumental efforts are being undertaken to increase the overall sensitivity [2, 3, 4, 5, 6], nevertheless, protein quantification remains a challenging task. In general, protein level intensities have to be inferred from peptide level intensities. This is complicated by the fact, that two peptides of the same protein - even though they are equally abundant in the sample - can be orders of magnitude different from each other in their measured intensities, for example due to differing ionization efficiencies [7] of the peptides. Additionally, ions with similar mass can interfere with the quantified peptides and distort the signal [1]. As many more low intensity than high intensity signals are present in a sample, interference of low intensity signals is common. A further challenge is due to *missing values* in the data. This denotes peptides that are only identified in some of the samples and missing in the other samples. Several setups are available for quantitative proteomics. In label free quantification (LFQ), each sample is measured in an individual liquid chromatography tandem mass spectrometry (LS-MS/MS) run and the peptide intensities are compared between the runs. In the most widely used setup, peptide intensities are derived from the full (MS1) scans [8]. The sets of peptides identified in each run are often not identical and, therefore, lead to missing values. This problem can be adressed by matching the MS1 peaks in the neighboring runs, but this solves the problem only to a limited extend.

A quantification approach that is less computationally challenging is chemical labeling via tandem mass tags (TMT) [9]. For TMT, up to 11 samples are isobarically labeled on the peptide level and mixed before submission to LC-MS/MS. The labels have reporter ions of distinct masses, which are detected in the fragmentation spectrum. Depending on the machine type, the fragmentation spectrum for reporter ion quantification can be a classical MS2 spectrum, or an intensity reduced MS3 spectrum, which is generated by further fragmentation of MS2 fragments [3].

In general, the challenges of protein inference, differing ionization, noisy peptide data and missing values are expressed to a certain degree in all quantitative MS setups and computational approaches have to deal with them appropriately.

A common approach for differential expression analysis is to derive protein level fold changes from the peptide fold changes and to apply statistical tests such as the t-test to assign a significance to it. This approach is for example implemented in the Perseus pipeline [10]. Peptide level models have been proposed [11, 12] and have recently been shown to offer superior performance compared to protein level approaches [13, 14, 15]. A recent implementation is given in the MSqRob package [14]. The majority of peptide level models are based on linear regression, which can be problematic for data with strong distortion, outliers, or small peptide numbers. We propose an orthogonal approach, consisting of the direct assessment of peptide level noise, which we term **M**ass **S**pectrometry analysis using **Empi**rical and **Re**plicate based statistics (MS-EmpiRe). We introduce empirically generated, intensity dependent error fold change distributions and utilize this for between-sample normalization and to derive differential expression probabilities for each peptide. We then show that these probabilities can be combined to the protein level via a modified Stouffer method [16]. The data for MS-EmpiRe can be measured with a variety of quantitative proteomics setups, as we need only peptide intensities grouped to proteins as input. We test the performance of the method on a recently published proteome wide benchmarking dataset of O’Connel, Gygi and coworkers [17], containing LFQ as well as TMT-MS3 data. With MS-EmpiRe we observe up to 121% more sensitivity in comparison to the approach reported in O’Connell et al. We additionally compare our approach with the peptide level tool MSqRob and see similar performance increases. On simulated differential expression changes, we see similar performance results as on the benchmarking set and demonstrate MS-EmpiRes superior classification abilities. MS-EmpiRe is available as a R package on GitHub (github.com/zimmerlab/MS-EmpiRe).

## 3 Methods

### 3.1 MS-EmpiRe compared to state-of-the-art approaches

We compare our method with two current methods, MaxLFQ with Perseus [8, 10] and MSqRob [13, 14], which have different strategies of solving the challenges associated with MS based protein quantification (Tab. 1). For the comparison, we focus on the principle steps of differential quantification: Normalization between different experimental samples, the statistical test applied, derivation of the corresponding test statistic (which represents the protein level change), estimation of the variance parameter(s) of the statistical test and outlier correction. In MaxLFQ, normalization is carried out by minimizing the sum of peptide level fold changes with run specific normalization factors. The sum is taken over all runs, also between conditions, with the underlying assumption that most of the proteins do not change. Several statistical tests can be applied to MaxLFQ data, where the t-test shows best performance [8]. The test statistic used in the t-test is the difference between the mean protein level LFQ intensities per condition. The LFQ intensities are pseudo-intensities derived from the median of the peptide level fold changes. The variance of the t-test is derived from the variance of the LFQ intensities between replicates. Outliers are implicitly taken care of by taking the median of the peptide level fold changes as a reference for the LFQ intensity.

**Table 1:** The two state-of-the-art differential quantification methods MaxLFQ (with t-test) and MSqRob compared against MS-EmpiRe.

|  | MS-Empire | MaxLFQ | MSqRob |
| --- | --- | --- | --- |
| Normalization | Median/Mode of peptide fold change distributions (only replicate samples) | Minimization of peptide fold changes (all samples) | Estimation of sample specific contribution via linear model |
| Fold change estimation | Maximum likelihood estimation from peptide fold change distribution | Fold change of MaxLFQ protein intensities | Estimation of condition specific contributions via linear model |
| Variance estimation | Empirical and Replicate based peptide distributions | Protein-level sample variance | Empirical Bayes Estimation |
| Outlier detection | Signal scoring based on outlier distribution, downscaling of outlier peptides | Median of peptide fold changes | M-Estimation with Huber weights |
| Statistical test | Modified Stouffer's method | t-test | Moderated t-test |
| Test statistic | Sum of Z-transformed, background normalized peptide fold changes | Difference of MaxLFQ protein intensities | Estimation of condition specific contributions via linear model |

The MSqRob method relies on a linear model, similar to the limma method for microarray data [18]. The linear model describes each peptide intensity as a composition of 4 different effects: The effect of the technical replicate, the effect of the biological replicate, the effect of the peptide specific ionization and the effect of protein level regulation. The normalization step is hence included in the linear model estimation. MSqRob uses the t-test as a statistical test. The test statistic is derived from the protein level regulation effect. Ridge regression and an empirical bayes appraoch are used to stabilize the intensity estimates and the protein level variance, respectively. To reduce the effects of outliers, M-Estimation with Huber weights is used, which shrinks the effect of high-residual observations [14]. MS-EmpiRe carries out normalization based on peptide level fold change distributions. Between-replicate distributions are used, which ensures that the fold change distribution should be centered around zero. The statistical test applied is a slightly modified version of the Stouffer method. This way, individual peptide level probabilities for regulation can contribute to the test statistic. The peptide level probabilities are derived from the peptide fold changes in the context of empirical, dataset-specific and intensity dependent background distributions. The variance estimation is hence carried out via these distributions. This allows a refined weighing of the influence of peptide noise on a given fold change. Outlier detection is carried out by explicitly modeling the influence of outlier signals and downscaling of outlier peptides.

### 3.2 Normalization

Mass spectrometry data suffer from sample specific effects, i.e. systematic perturbations which affect whole samples. For instance, the total amount of protein which is processed per run has a significant effect on the signal measured per peptide. Raw signals originating from two different samples are therefore hardly comparable. To correct for sample specific effects, all signals of a sample are typically multiplied by a *scaling factor*. In the context of RNA sequencing data, which are subject to sample specific effects as well, procedures to detect appropriate scaling factors are introduced e.g. by DESeq and edgeR [19, 20, 21]. While DESeq finds scaling factors by comparison of every sample to a virtual sample, edgeR computes all pairwise scaling factors. Both methods use the median of many gene level fold changes as an estimate for the scaling factor. MaxLFQ [8] is a normalization procedure for mass spectrometry data. Instead of relying on the median, MaxLFQ solves a system of linear equations to identify scaling factors such that the change of peptide signals between any two samples (and fractions) is minimized.

All previously mentioned normalization procedures rely on the assumption that most of the signals do not change between any two samples, even when samples from different experimental conditions are compared. We propose to reduce this assumption to samples from the same experimental condition (i.e. replicate samples) and use a different factor to normalize between conditions.

Normalization within a condition in MS-EmpiRe is done by single linkage clustering as described in Fig. 1. Each cluster contains either one or multiple samples. We start with as many clusters as we have replicates and successively merge the two most similar clusters until we end up with one cluster that contains all samples. Similarity between any two clusters is defined as follows: For two samples, we compute a *fold change distribution*. We build every possible sample pair between the two clusters and compute the fold change for every peptide which was detected in both samples of a pair. The variance of this distribution is used to determine the similarity between clusters while the median is used as an estimate for a systematic signal shift. To merge two clusters *c*_1_ and *c*_2_ we scale all signals of samples in *c*_2_ by the median. This step yields a new cluster that contains all samples from *c*_1_ and *c*_2_ and in which all samples are shifted onto each other.

**Figure 1:**
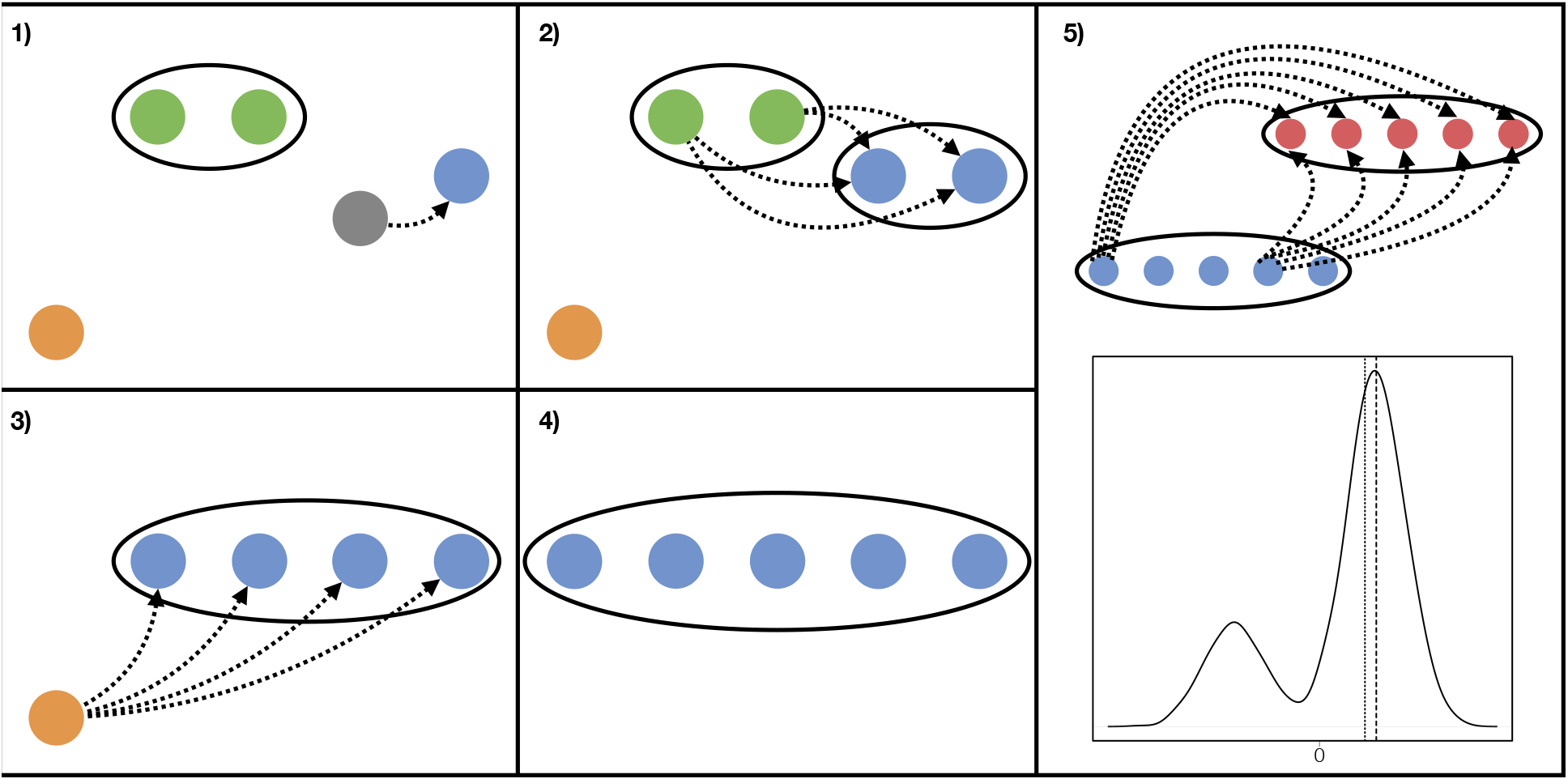
Single linkage clustering for signal normalization. **1)** We start with one cluster containing two samples (green) and three clusters of size one. We identify the two nearest clusters (grey and blue) and merged them to one new cluster by shifting the signals of the grey sample according to the median log_2_ fold change to the blue sample. **2)** We merge the green and blue cluster. Since they both consist of multiple samples, we determine the shift parameter by computing signal fold changes between any possible inter-cluster sample pair. **3)** The last cluster merge step **4)** The algorithm results in one cluster containing all samples. **5)** The two final clusters of different conditions are merged (blue and red). The resulting distribution of all inter-cluster fold changes is shown below.

Single linkage clustering is applied to each condition separately. Samples from two different conditions are then shifted in a similar way, the difference is the selection of the shift parameter. Since we can no longer assume that none of the peptides changes, we propose to use the most probable fold changed from the distribution instead of the median. This choice similar to the idea of centralization proposed by Zien et al. [22]. Instead of enforcing a minimal change between all peptides, this shift only targets the majority of peptides. and is still in accordance with the assumption that most proteins do not change. The shift parameter can also be defined by the user using prior knowledge for certain subsets of proteins for example.

After the normalization, signals for the same peptide between any two samples are comparable.

### 3.3 Empirical error distributions

Our goal is to detect differentially abundant proteins between different conditions. However, only peptide level measurements are available from current standard MS experiments and protein level changes have to be inferred. We argue, that each peptide level change should be assessed in context of the noise associated with the measurement. MS-EmpiRe is therefore centered around replicate based *empirical error distributions* (Fig. 2.1 and 2.2). The empirical error distribution is fully based on the data and derived as follows: We compute the log_2_ fold change of every peptide signal between any two replicate pairs in each condition. As the log_2_ fold change between replicate samples should be zero, each deviation from zero can be seen as an error. This results in one large collection of errors, approximately 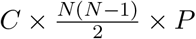 for *C* conditions with *N* replicates each and *P* detected peptides. Since we observed that the variance of peptide measurements depends on signal strength (Fig. 3 e) we decided to split the complete distribution into intensity dependent sub-distributions. Each of the resulting sub-distributions contains just a subset of all peptide fold changes. For the construction we sort the peptides ascending to their mean signal strength. We slide a window over the sorted list of peptides to determine the relevant subset for each distribution. The window size and how far it is shifted in each step are parameters that can be controlled by the user. Adjusting it can increase the resolution of the sub-distributions at the cost of computational time. The default window size is 10% of the total number of peptide measurements with a maximum of 1200. The size of the shift is set to 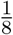 of the actual window size. Note that each peptide may appear in multiple sub distributions if the shift is smaller than the window size. To assign each peptide to only one sub-distribution, we save the mean signal of the first and last element of each distribution. We then calculate the distance of the mean signal to the start and end of each distribution for every peptide. Each peptide is then assigned to a distribution such that the minimum of those two distances is maximized, i.e.

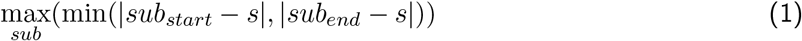

After this step we have a collection of empirical error distributions that describe the observed measurement errors in relation to signals.

**Figure 2:**
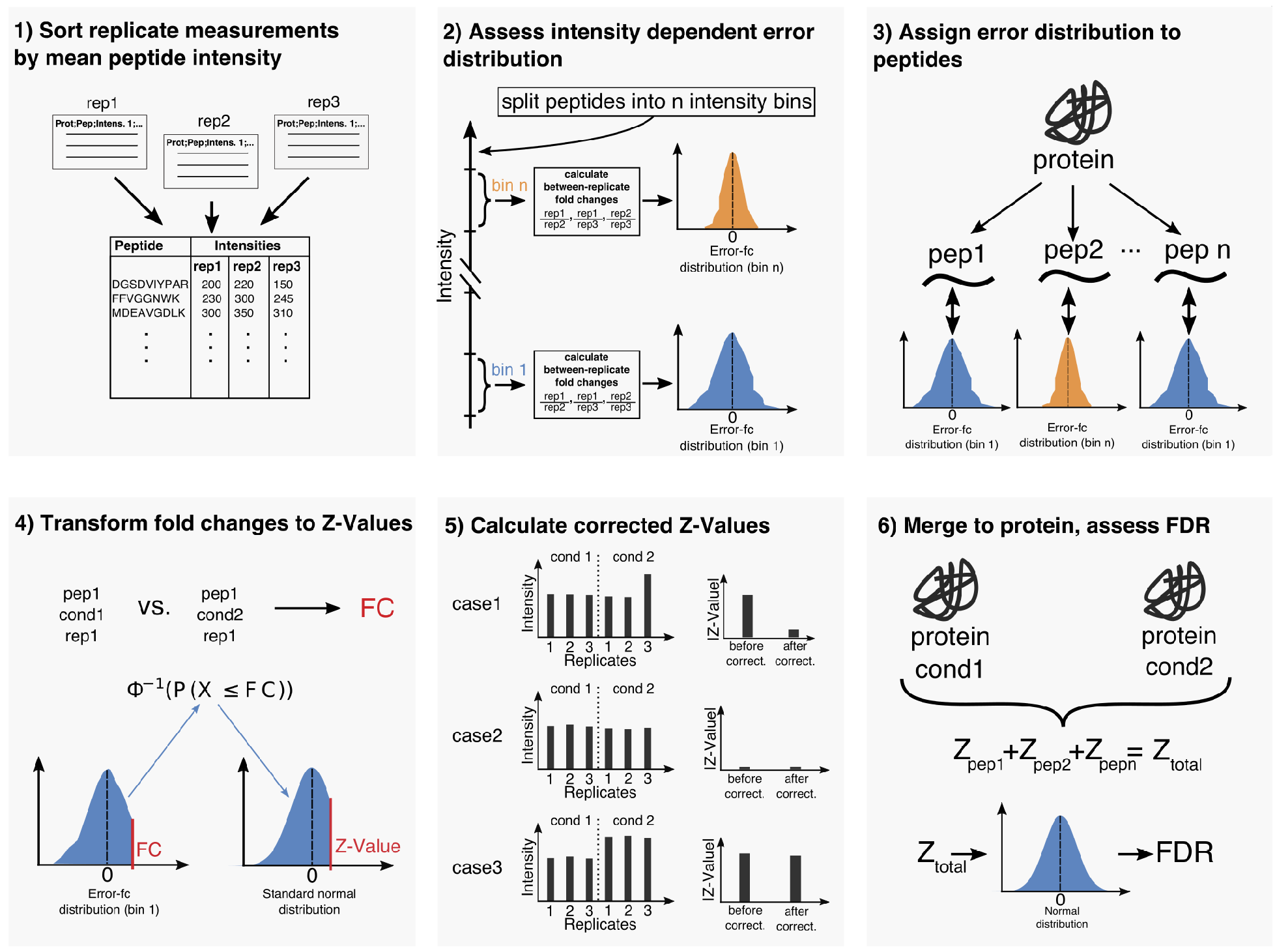
Schematic of the MS-EmpiRe workflow. 1) All identified peptides from a proteomics run are sorted by their mean intensities. 2) The peptides are the split into subgroups based on their intensity. For each subgroup, the *error fold changes* of the individual peptides are calculated. An error fold change simply denotes the *log*_2_ fold change of a peptide between two replicate conditions. All error fold changes within a subgroup form an *empirical error distribution*. Distributions corresponding to lower intensity peptides show a larger variance than for high intensity peptides. 3) When a protein is tested for differential quantification, each peptide gets assigned an empirical error distribution. Peptides of similar intensities can get the same distribution assigned. 4) For each peptide fold change, the probability that this fold change happened by chance (e.g. the p-value) is assessed from the empirical distribution. This means that the same fold change will get a much lower p-value when the distribution is wide as compared to when it is narrow. To make this value manageable, the p-value is then transformed to a Z-value, by transferring the mass of the empirical probability distribution to a standard normal distribution. 5) The Z-values for each peptide are corrected for outliers. For this, the probability is estimated that a high Z-value on the peptide level has happened by chance due to individual outliers. 6) The corrected Z-values can directly be summed to the protein level and the corresponding protein-level FDR can be obtained after multiple testing correction.

**Figure 3:**
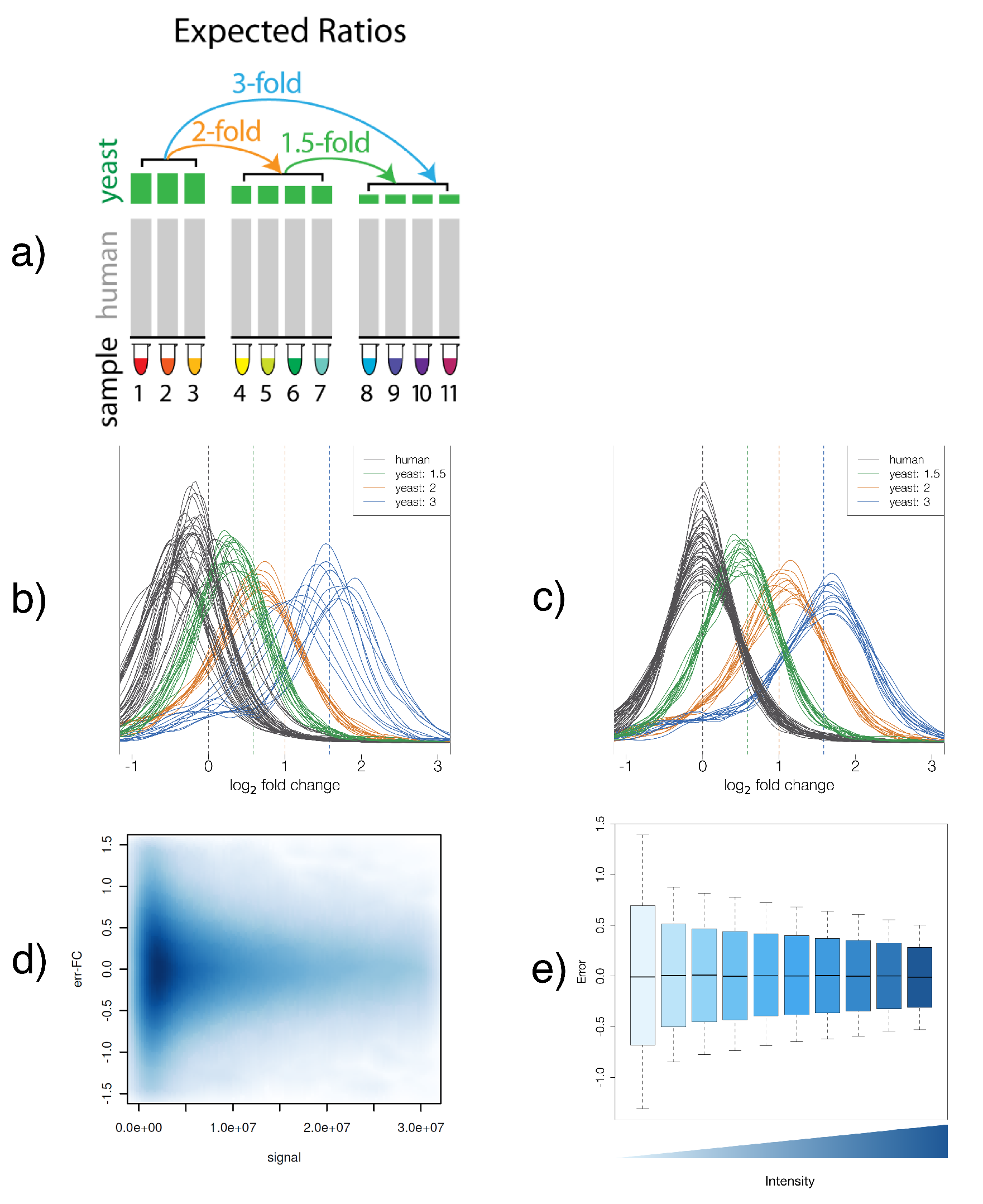
Experimental setup and fold change based metrics. a) Benchmarking setup for quantitative assessment of fold changes taken from OConnell et al.[17]. Different amounts of yeast lysate are spiked into human cell lysate. The three groups contain 10%, 5% and 3.3% yeast lysate, respectively, b) Peptide level fold change distribution between all conditions, before normalization. c) The distribution after fold change based normalization. e) Intensity dependent peptide fold changes between two replicates of the LFQ data (error fold changes), displayed as smoothed density scatter plot. f) Error fold changes for 10 intensity regions displayed as box plots. Each box contains the same number of data points. The quantiles correspond to the fractions 0.05, 0.15, 0.50, 0.85, 0.95.

Any observed peptide fold change can now be put into context of the background noise. This allows to determine the probability of the peptide fold change under the corresponding empirical error distribution. We denote this probability as the *empirical p-value*.

### 3.4 Merging scores over replicates

We can now determine the empirical *p*-value for every peptide between any two samples. What we rather want, however, is the same information for whole proteins between two conditions including replicate data. This means we have to express the empirical *p*-value in terms of a score that we can combine over replicates as well as peptides. Furthermore the score should be able to distinguish between negative and positive fold changes. This way we can identify groups of peptides that consistently show the same direction of change between multiple replicate pairs. One score fulfilling these criteria is the *Z*-value, i.e. a score that follows a standard normal distribution. We can transform an observed fold change into the corresponding *Z*-score as follows:

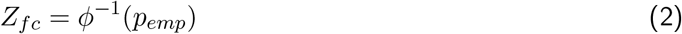

where *ϕ^−1^* is the inverse of the cumulative distribution function of the standard normal distribution and *Pemp* is the empirical *p*-value. This is analogous to Stouffer’s method [16] for combined probability tests.

This means we can transform any empirical error distribution to a standard normal distribution (Fig. 2.4). In the following sections we will show how those *Z*-scores can be transformed to joint probabilities over replicate data as well as multiple peptides.

To distinguish between background noise and signals, usually not only 2 samples, but *N* vs *M* replicate measurements from two different conditions are compared. Those yield up to *N* × *M* scores per peptide which are merged to make a protein-level statement between the two conditions. Under the null hypothesis of no change, each of the *N* × *M Z*-values follows a standard normal distribution. Under this assumption, we can simply compute the sum of the *N* × *M* standard normally distributed *Z*-scores which follows a normal distribution as well. Looking at the sum is reasonable for the following reasons: It should become extreme only if we have multiple measurements that consistently deviate in the same direction. Too few too weak deviations are canceled out by the non deviating measurements. The same is true for strong deviations in different directions. The mean of the resulting normal distribution is zero as it is the sum over the individual means. In general, the variance of a random variable that is the sum over multiple random variables is the sum over the full covariance matrix of the variables, i.e.

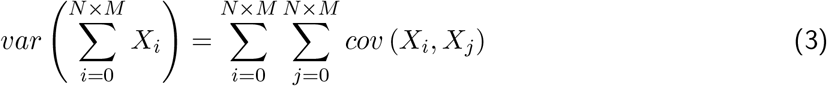

The means and variances are known for each of the variables since they follow a standard normal distribution. We are also able to compute the covariances for dependent variables. This is necessary because some of the possible sample comparison are not independent, in particular any two sample pairs that share either the first or second sample. It can be shown that the covariance of two *Z*-scores random variables that share one of the samples is 0.5 (Supplementary Material, Section 1).

For each peptide, we can now unexpectedness under the previously derived background distribution over all sample pairs.

### 3.5 Correcting for outlier measurements

One problem about the sum described in the previous section, is that it is susceptible to single outlier measurements. A single extreme *Z*-score can be sufficient to make the resulting sum significant. This is because of the null hypothesis that *each* of the sample pair comparisons must not be differential. We therefore introduce a correction to estimate the probability, that a single outlier shifts the distribution towards higher values (Fig.2.5). For this correction, we estimate the *Z*-value of the peptide when it is not regulated (*Z_normed_*)and substract it from the original *Z*-value (*Z_orig_*). *Z_normed_* is estimated as follows: We compute all possible fold changes of the peptide between two conditions (replicate 1 vs. replicate 1, replicate 1 vs. replicate 2, etc.). This results in a (very small) fold change distribution. Analogous to section 3.2, we use the median of this distribution as a scaling factor and shift all signals of the second condition by the median. This minimizes the difference of signals between the two conditions and simulates a non-regulated peptide. We again compute the summed *Z*-value for those shifted peptides, i e *Z_normed_*. If the peptide measurements were differentially regulated previous to the shift, *Z_normed_* would be less extreme than *Z_orig_*. If the shift does not change the signal, *Z_orig_* and *Z_normed_* are more or less the same. We can hence introduce a new value *Z_corrected_ = Z_orig_ — Z_normed_*, which denotes the difference between a regulated and a non-regulated shift. We now want to use the distribution of *Z_corrected_* to estimate, how unlikely an observed *Z_corrected_* value is. The higher *abs(Z_corrected_)* is, the more extreme the original measurement was.

However there exists no closed form for the distribution of *Z_corrected_*. We therefore sample such a distribution by simulating a set of non-differential measurements. For each simulated measurement, we compute *Z_corrected_*. Similar to Eq. 2, we look up the cumulative probability of a measured *Z_corrected_* in the simulated distribution of *Z_corrected_* and transform its empirical *p*-value to a *Z*-value. Whether this correction is needed may depend on the data. After the correction we end up with a score which is standard normally distributed under the null hypothesis of no change.

### 3.6 Correcting for outlier peptides

The previous section shows how to correct for outlier measurements of single intensities. Peptides that show a much more extreme change over many samples than the remaining peptides of the same protein have to be accounted for separately. To detect outlier peptides, we compute fold changes for every peptide and sample pair of the same protein. If the median of a single peptide is more extreme than the 25%/75% quantile of the whole protein distribution and protein and peptide are shifted in the same direction, it is marked as an outlier.

To compute *Z_normed_* for an outlier peptide, we shift the signals by the 25%/75% quantile of the protein distribution instead of the median of the peptide distribution. This modified shift results in a less extreme *Z_corrected_* for the peptide.

### 3.7 Combining the peptide scores

What we have so far is a *Z*-score that expresses how likely it is for a peptide fold changes over all possible replicate comparisons to occur by chance. Each of those *Z*-scores follows a standard normal distribution.

Similar to the first step of merging the peptide scores over all replicate pairs, we can join those scores for all peptides from the same protein (Fig. 2.6). In contrast to measurements for the same peptide from different sample pairs, peptide measurements can be regarded as independent measurements for the same protein. This means that under the null hypothesis that every peptide score is a standard normal distributed variable, the sum of such peptide scores is distributed

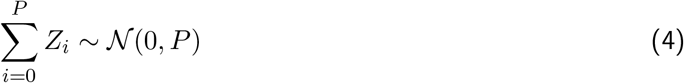

with *P* being the number of different peptides mapped to a certain protein. Using this sum of peptide score we can now express the probability of a protein under the null hypothesis of no change while taking into account all replicate measurements. To correct for multiple testing we finally apply the Benjamini-Hochberg false discovery adjustment.

### 3.8 Re-processing of the proteome wide benchmarking dataset

We downloaded the raw data of the study of OConnell et al. [17] from the PRIDE repository PXD007683 and processed the TMT as well as the LFQ dataset with MaxQuant [23] version 1.6.0.16 with standard settings and the respective quantification set (11 plex TMT-MS3 and LFQ). Each set contained three conditions: 10% yeast spike-in, 5% yeast spike-in, 3.3% yeast spike-in. For 10% yeast spike-in, three replicates were measured, for 5% and 3.3%, four replicates were measured. The datasets were searched against a combined yeast (7904 entries) and human (20317 entries) database downloaded from Uniprot (04/2018, reviewed). Specific digestion was set for trypsin with two missed cleavages allowed. Carbamidomethylation of Cysteine was set as a fixed modification and Oxidation of Methionine and N-terminus Acetlyation were set as variable modifications. 20ppm mass tolerance were set for precursor ions und 0.5m/z were set for fragment ions. Results were filtered to 1% FDR at the peptide-spectrum-match (PSM) and protein level. For the LFQ data, the match between runs option was set (default configuration). We compared the number of identified proteins with the results reported from O’Connell et al. and observed only slight differences, possibly due to different databases and MaxQuant versions (−2% for TMT and −1% for LFQ for our protein). Hence the numbers are a bit different but should not influence the overall outcome. We found around 400 proteins after filtering in the Perseus/MaxLFQ setup, which agrees well with O’Connell et al. As we used MaxQuant instead of SEQUEST [24] for the TMT analysis, we had a noticeably lower identification rate (around 100 proteins less compared to the SEQUEST results presented in the main text of O’Connell et al.). This is similar to the number of MaxQuant results for TMT, reported in the supplement of O’Connell et al.

### 3.9 Filtering of the benchmarking dataset

Between the different tools, we noticed large differences in the number of proteins that are actually submitted to statistical testing. MaxLFQ with Perseus showed the most conservative filtering, while MSqRob was most permissive. Most impact on the filtering had the decision, whether to also accept proteins with only one quantified peptide value. With only one peptide per protein, a misidentified peptide can immediately lead to a false classification. As in MS-EmpiRe, a peptide needs to be consistently quantified over multiple replicates to gain significance, the probability for such an event decreases and we hence decided to also use a less conservative filtering of only one peptide per protein. We also compared the one-peptide with the two-peptide approach and saw no significant effects on the FDR (see supplemental Fig. 5). This underlines that MS-EmpiRe is designed appropriately deal with sparse peptide evidence caused by many missing values. For filtering of MS-EmpiRe the following peptides/proteins were excluded: reverse peptides and contaminants, peptides mapping to yeast as well as to human and proteins quantified in only one replicate.

### 3.10 In silico benchmarking

The HeLa background proteins from the study of O’Connell et al. were normalized via MS-EmpiRe and each sample was considered a replicate measurment. This resulted in 11 quasi-replicate runs, out ouf which 6 were randomly chosen. The 6 replicate measurements were split into two sets with 3 replicates each. One of the sets was chosen for in silico expression changes. For the selected set, a subset of the proteome was chosen and was artificially ‘‘regulated”. For each protein in the subset, an expression change factor was drawn from a distribution. The peptide level changes for the protein change were then sampled around this factor. The changed and the unchanged subset were then compared as two separate experiments with MS-EmpiRe. As in the benchmark it was known which proteins were regulated, the differential quantification performance (sensitivity, specificity etc.) could be assessed.

## 4 Results and Discussion

### 4.1 Fold change based normalization reveals structure of the benchmarking dataset

We used the benchmarking dataset from the study of O’Connell et al. [17], where yeast is spiked into human cell lysate at different concentrations (10%, 5% and 3.3%) (figure 3a). Hence, when comparing the abundance of yeast proteins e.g. in the 10% sample with the 5% sample, one expects a fold change of 2 for each yeast protein. The two other combinations 5% vs. 3.3% and 10% vs. 3.3% give a fold change of 1.5 and 3, respectively. When applying a differential quantification algorithm, the changing proteins are known (e.g. the yeast proteins) and, thus, measures like specificity and sensitivity can be assessed. The samples were measured twice, once using a TMT-MS3 approach and once using label free quantification (LFQ). To visualize the normalization procedure employed by MS-EmpiRe, we used the LFQ dataset. Between every sample pair, we calculated the *log_2_* fold change for each individual peptide. This resulted in a distribution of fold changes for each sample pair, which was either a between replicate, or a between sample distribution. For the between replicate distribution, we would only expect deviations from 0 due to measurement errors or biological variation and, therefore, we call this distribution the *empirical error distribution*. For the between sample distribution, we would only expect systematic deviations from 0 for the regulated proteins. In Fig. 3b, the between sample distributions are displayed before normalization. For clarity, human proteins (which should not change at all) and yeast proteins (which have systematic changes applied to them) are displayed separately. We can already see some trends in the distributions, which underlines the fact that the fold change based view is an intuitive measure for quantitative datasets. When applying subsequent between-replicate and between-sample normalization, as described in the methods section, we obtain the visibly clustered distributions displayed in Fig. 3c. The human peptides (around 90% of peptides) are not shifted and distributed around 0. The yeast proteins are aligned around the shift that was experimentally applied to them (i.e. the *log_2_* transformation of 1.5, 2 or 3). If too much or too little yeast had been applied to one of the samples, this would reflect in a stronger deviation in a subset of the replicate distributions. This is not the case in our dataset and we see with slight deviations-an alignment around the desired value (dashed lines). In a real life example, we would not expect systematic changes around one fold change in one direction, but larger spread deviations in both directions. The example, however, visualizes that a fold change based approach on quantitative data sets is an effective procedure to normalize datasets without altering the structure of the underlying data. In general, the distributions reveal a ubiquitous problem in MS based proteomics data. The data is so noisy, that a lot of the measured yeast peptides do not even show regulation. This is most striking for the 1.5 set, where around one quarter of the peptides show no regulation, or even regulation into the wrong direction. Hence and there is no way to classify these peptides correctly by themselves. Since usually, multiple replicates and peptides exist for a protein, the quantification of a protein can be seen as multiple drawings from such a distribution. This underlines that peptide fold changes should always be analyzed in the context of the dataset specific noise.

### 4.2 Assessment of empirical error distributions underlines the importance of context dependent fold changes

As already described in the methods section, one of the key features of the MS-EmpiRe algorithm is the quantitative assessment (and subsequent utilization) of fold change consistency. For this we make use of the fact that the log fold change between replicate measurements should be 0, which allows us to derive empirical error distributions containing the fold changes of peptides between replicate measurements. These distributions have already been discussed in the previous section and the human peptides displayed in Fig. 3c are a good example. However, when looking at the empirical error distributions over a whole dataset, as in the previous section, we neglect the well-known fact that low-intensity peptides are subject to significantly more variation than high intensity peptides. An intuitive way to visualize this is by plotting the error fold changes against the mean intensities of the peptides, as displayed in Fig. 3d). In this density scatter plot, we see that the majority of peptides is actually of low intensity and in the low intensity region a large spread of the fold changes is visible. Furthermore, we see global dependence of the error fold changes on the intensity and high intensity peptides are much less prone to deviate strongly from 0. Nevertheless, outliers exist for all intensities, which underlines the need for a more quantitative assessment. This is depicted in Fig. 3e), where empirical error distributions are given for distinct intensity bins. The original fold change distribution is split into 10 boxes and each box contains equal amounts of data points. We see that the lowest intensity box is particularly noisy, with around 40% of the data above a fold change of 1.5 (*log_2_* fold change of around 0.6). Hence, if a low intensity peptide has a 50% intensity increase from one condition to the other, the likelihood that it is an actually irrelevant change is at around 40%. When we consider the highest intensity box, this value drops from 40% to around 5%. Identical fold changes hence have a very different meaning, depending on the context. The MS-EmpiRe approach transforms an observed fold change in the context of its corresponding empirical error distribution and, therefore, quantitatively accounts for this phenomenon. Especially, as every datasets carries its own noise, and e.g. TMT-MS3 data shows significantly reduced noise (see supplemental Fig. 4) the empirical assessment for each dataset is crucial.

**Figure 4:**
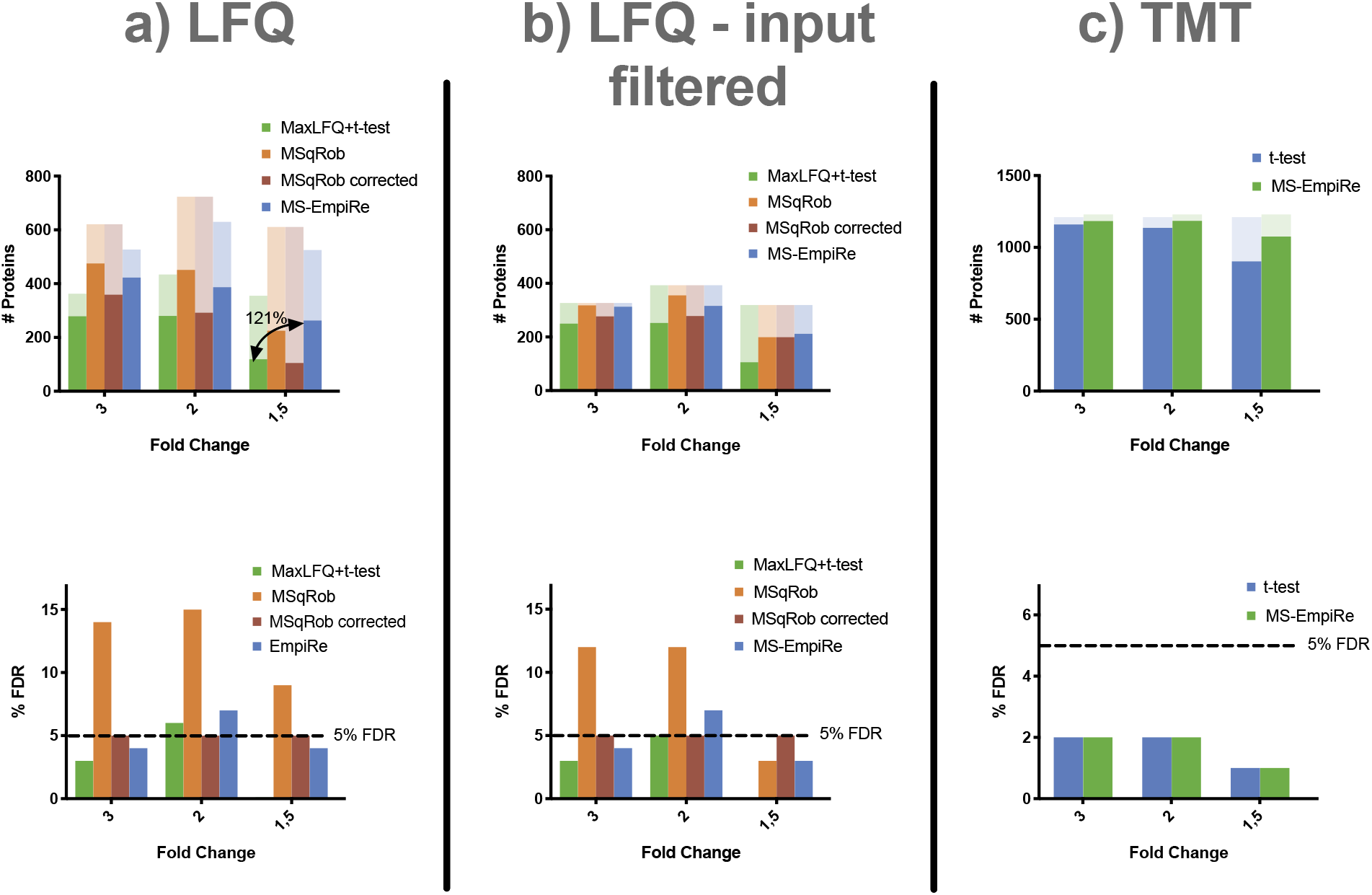
Assessment of the differential detection performance on the benchmarking setup of OConnell et al. a) Number of proteins detected in the LFQ data by the Perseus setup, MSqRob and MS-EmpiRe. The light shades show the numbers of yeast proteins accessible for testing in the setups, which differ for every method. As MSqRob shows high FDR rates (bottom plot), an FDR-corrected bar is introduced for MSqRob. Perseus shows low sensitivity at good error rate control similar to MSqRob. Perseus is very conservative for the fold change 1.5 setup, with no false positives (no bar visible). MS-EmpiRe increases the detection substantially in all cases with good error rate control. This is most pronounced in the most challenging fold change 1.5 setup. b) Number of proteins detected when using an intersected input set. Due to this conservative approach the numbers and differences are lower in general, nevertheless MS-EmpiRe is the best performing method. c) Comparison of Perseus and MS-EmpiRe on a TMT dataset. MSqRob was excluded as it currently does not support TMT data. The overall performance is better due to higher depth from sample fractionation, lower noise and fewer missing values. Quantification on the protein level hence already works well, still MS-Empire shows a significant sensitivity increase of around 19% for fold change 1.5.

### 4.3 MS-EmpiRe shows up to 121% sensitivity increase compared to state of the art approaches

The key question addressed in the setup of O’Connell et al. [17] is, how many proteins can be detected as differentially expressed in a proteome wide benchmarking setup. Analogous to their paper, we assessed how many of the experimentally shifted yeast proteins we were able to detect via MaxLFQ coupled to the Perseus pipeline. We then compared this approach with our MS-EmpiRe approach and the recently published tool MSqRob. Perseus was executed analogous to the settings given in O’Connell et al. (reverse and contaminant filtering, at least two replicate measurements per protein and two sided homeoscedatic t-test with Benjamini-Hochberg correction) and MSqRob was executed with default settings. A FDR of 5% was set for all approaches. In Fig. 4a) and 4b), we show the results of the benchmark for the more challenging LFQ setup. The number of peptides available for testing differs markedly, depending on how conservative the corresponding tool is in filtering peptides for quantification (see also methods section). MS-EmpiRe clearly outperforms Perseus in terms of sensitivity, with up to 120% more differential protein detections in the fold change 1.5 setup. When comparing MS-EmpiRe with MSqRob, it seems that MSqRob is slightly more sensitive. However, for MSqRob the observed FDR (i.e. the number of human proteins detected) is between 9% to 15% instead of the required 5% for MSqRob. Perseus and MS-EmpiRe also violate the FDR in the fold change 2 setup, but only by 1 and 2 percentage points, respectively. To make the sensitivity analysis more comparable, we show an FDR corrected bar for MSqRob, where we set the FDR cutoff of MSqRob to a more stringent value (see methods section), such that the actual FDR is also at around 5%. In this setup MS-EmpiRe outperforms MSqRob in terms of sensitivity in all cases and detects more than twice the number of proteins of MSqRob in the most challenging fold change 1.5 setup. As MS-EmpiRe and Perseus both violated the FDR for the fold change 2 dataset, we looked at the corresponding data in more detail. We saw that many of the misclassified human proteins in the dataset were particularly tough cases, where many peptides over many replicate conditions show consistent up- or down regulation (see supplemental table 1). Of course, cases like this are included in the FDR estimation. However, the condition seems to show over-proportionally many cases of consistent protein up-regulation. We considered tuning the FDR estimator to be more conservative, but took into account that the FDR violation was only mild and that neither the other benchmarking conditions, nor the simulations showed further FDR violations. As slight regulation or systematic distortions might always occur under experimental settings, we decided to leave the model as is. The setup shown in Fig. 4a contains the input sets after the individual filtering applied by each method. This corresponds to a real-life application of the methods, but also reduces the comparability of the classification capabilities. We hence compared each method on the same set of peptides, which consisted of the intersection of all input peptides. This led to a significant decrease of the number of proteins detected. Interestingly, the drastic reduction of input peptides strongly increased the number of detected proteins for MSqRob for the 1.5 set. This implies, that MSqRob is prone to give an over-optimistic scoring to proteins with sparse peptide evidence and hence more stringent filtering might be appropriate. In the fold change 1.5 set, MSqRob also does not violate the FDR constraint. In the two other sets, the peptide filtering does not seem to suffice to control the FDR and also for MS-EmpiRe the fold change 2 set still violates the FDR slightly. Nevertheless, MS-EmpiRe is the most sensitive method over all sets. When comparing the methods on the less challenging TMT data set in Fig. 4c), we see overall high sensitivity, which is also discussed in O’Connell et al. Also here, we see a substantial increase in sensitivity of around 20% compared to Perseus, when considering the 1.5 fold change set.

### 4.4 In silico benchmarking shows high sensitivity and conservative FDR estimation

The importance of experimental benchmarking setups for quantitative proteomics cannot be overstated. Without reference standards, it is impossible to estimate the performance of experimental and computational methods. Unfortunately, performing an experimental benchmarking is cumbersome as it requires very precise mixing and sample handling. Additionally, only constant fold changes can be applied to a given setup, which does not reflect an actual regulative scenario. To complement the experimental benchmarking, we generated an in silico benchmarking set, as described in the methods section. In short, we used the human background proteins measured in O’Connell et al. as replicate measurements and we divided six replicate measurements into two groups. We then applied *in silico* intensity changes on the protein and peptide level to one of the groups and compared the two groups in a differential quantification context. As we know which proteins are ‘‘artificially regulated”, we can assess measures like sensitivity and specificity analogous to the experimental benchmarking setup discussed in the previous sections. We simulated two setups: one similar to the one in O’Connell et al., where we always applied the same fold change (with some noise) to a sub-fraction of the proteome, including a 10% fraction. Additionally, we simulated a more realistic scenario, where the *in silico* expression changes were not always the same, but were drawn from a distribution. We designed the distribution to be bimodal such that up- and down regulation was possible. The results for sensitivity and precision (i.e. specificity) for LFQ data are depicted in Fig. 5. The boxes result from changing different fractions of the proteome (individual simulations where 5%, 10%, 40% of the proteome are changed). Surprisingly, we noticed that the fraction of proteome changing significantly influences the sensitivity of the applied statistical test, especially for the t-test applied with Perseus. For example, when 30% of the proteome is changing with a fold change of 1.5, this is better detected by a statistical test as when only 5% of the proteome is changing with a fold change of 1.5. The reason for this is apparently a loss of significance after multiple testing correction. As multiple testing correction can be seen as a shifting of the p values into the direction of a uniform distribution, stronger deviations from the uniform distribution (e.g. many regulated proteins) are less strongly affected. In supplemental Fig. 6 we see, that the protein level scoring underlying the t-test does not allow a very distinct discrimination between regulated and non-regulated proteins as compared to MS-EmpiRe, which explains the losses in sensitivity with Perseus. The clearer distinction between regulated and non-regulated proteins by the peptide level tools is also reflected in the fact, that the peptide level tools MS-EmpiRe and MSqRob show less dependence on the proteome fraction in terms of sensitivity. In general, the results of the in silico simulations in Fig. 5 show a similar picture as compared to the experimental benchmarking setup. MS-EmpiRe and Perseus show conservative error estimation, which however comes at the cost of drastically reduced sensitivity for Perseus. MS-EmpiRe is the most sensitive tool and MSqRob detects only slightly less proteins. However, MSqRob shows problems in terms of error rate control, especially for setups with strong fold changes. This might be due to an over-optimistic error estimation of MSqRob due to the many clear classification cases. Comparing the fixed setup with the setup where we generate dynamic noise, we see that the overall identification rate decreases, whereas the general trends for all three methods are very similar. Error rate estimation does not decrease and hence all methods show the desired response towards high noise in the data.

**Figure 5:**
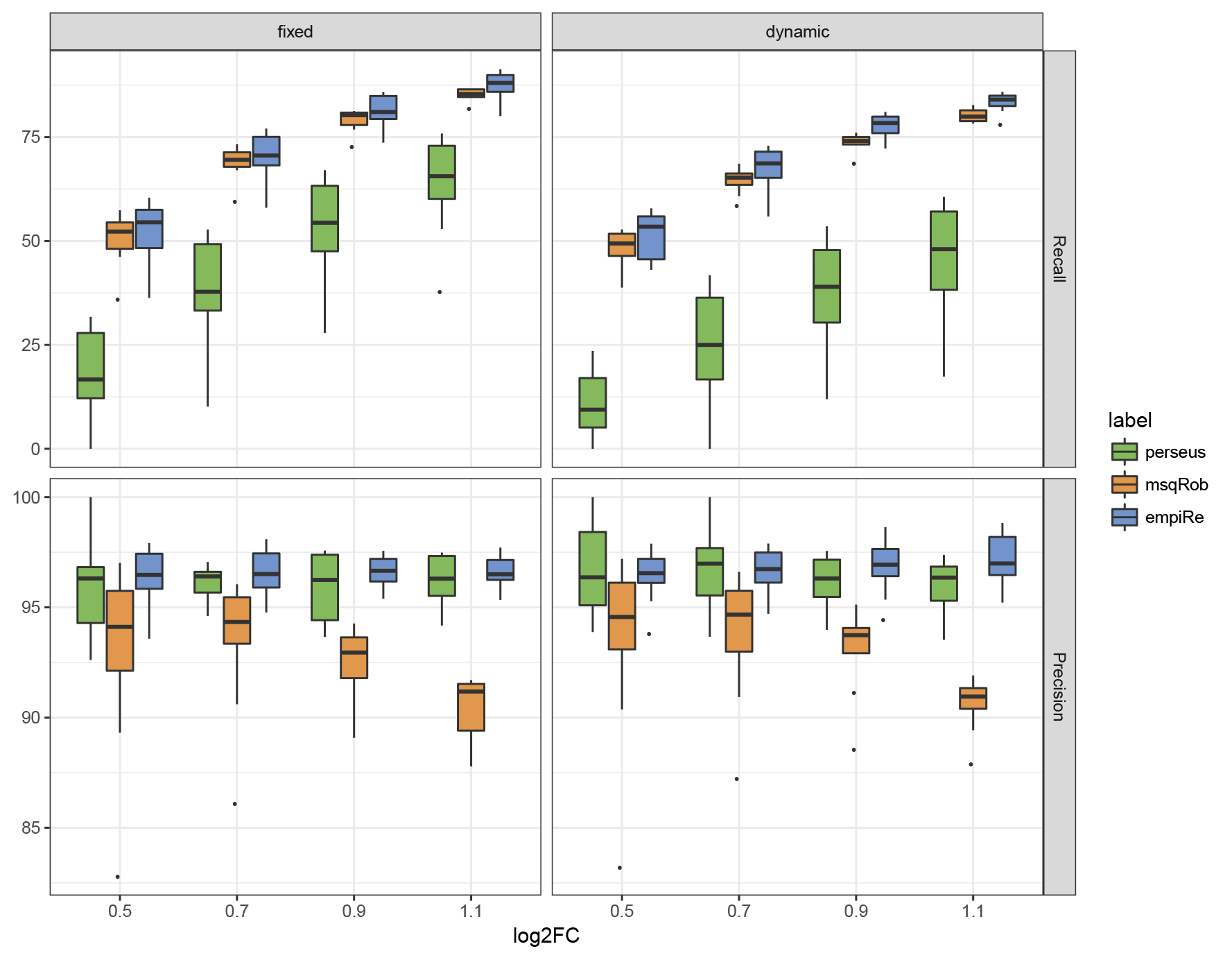
In silico benchmarking of MS-EmpiRe, Perseus and MSqRob. The x tics represent the median fold change by which the data is shifted. The boxes contain the sensitivity/specificity results, when different fractions of the proteome are changed. In particular each box contains eight values corresponding to 5% up to 40% of the proteome changing in 5% steps. Sensitivity and specificity for different fold changes upon constant (left) as well as dynamic (right) proteome changes are shown. A clear dependency on the fraction of the proteome changing is visible. As in the benchmarking set, Perseus shows low sensitivity with good error rate control, MSqRob shows high sensitivity but often violates the error estimation and MS-EmpiRe shows high sensitivity with good error rate control.

## 5 Conclusion

Current Mass Spectrometry proteomics publications often report the number of quantified proteins for a given proteomics setup. This number however, can be misleading, as the number of quantified proteins does not necessarily reflect the number of proteins that can actually be detected in a differential quantification experiment [17]. This is especially the case for more noise-prone proteomics setups, such as LFQ. Given the popularity of such setups, it becomes evident that increasing the depth of a proteomics at the level of differential detection becomes an ever more important aspect. Considering recent studies [13, 14], it is likely that the future development of differential quantification will go beyond the protein level, which neglects valuable information. Peptide level tools like MSqRob show significantly improved sensitivity, though we have shown that FDR control is still difficult in a proteome-wide setup. The popular MaxLFQ+Perseus pipeline also implements the most conservative approach in the experimental benchmarking setup. With MS-EmpiRe, we introduce a new peptide level tool that shows high sensitivity at accurate error rate estimation. While FDR estimation for MS-EmpiRe is almost equal to Perseus in the experimental setup, it even outperforms Perseus in the in silico simulation. We have shown that MS-EmpiRe gives up to two fold increase in sensitivity for small fold changes (1.5 fold change), which are highly relevant for biological applications. Even though a fold change of 1.5 is already very challenging for a proteomics setup, such a change may already reflect a drastic alteration in a biological system. The key to the sensitivity of MS-EmpiRe is the direct modeling of errors on the peptide fold change level. This gives an immediate statistical weight to individual peptide fold changes, which are then transferred to the protein level via basic statistics and without additional optimizations or parameters. This simple approach is enabled by the assumption of consistency between replicates and, therefore, our method heavily relies on replicate measurements. Even though MS-EmpiRe is able to process only two replicate measurements per condition, ideally three or more replicate samples should be available. For MS proteomics data, where robust workflows exist and the creation of replicate measurements is a standard, we believe that this requirement matches well with the current experimental practices. From our perspective, the consistency of replicates is a minimalistic and reasonable assumption that can be made for proper processing of proteomics data. A possible deviation from replicate consistency might occur, when uncontrolled factors in an biological experiment change between replicates. In our setup, this might lead to an underestimation of differentially expressed proteins. However, replicate-inconsistent setups are highly critical and should be handled with care. Based on our analysis, we conclude that MS-EmpiRe is currently the most sensitive tool for differential protein detection. MS-EmpiRe requires as inputs only peptide intensities and protein identifications and is therefore applicable to virtually any modern proteomics measurement. We hope that our tool becomes an easy-to-use option for proteomics researchers and helps to improve the quality and biological insight gained from MS proteomics studies.

## References

[1] Marcus Bantscheff, Simone Lemeer, Mikhail M. Savitski, and Bernhard Kuster. Quantitative mass spectrometry in proteomics: Critical review update from 2007 to the present. Analytical and Bioanalytical Chemistry, 404(4):939–965, 2012.

[2] J. V. Olsen. Parts per Million Mass Accuracy on an Orbitrap Mass Spectrometer via Lock Mass Injection into a C-trap. Molecular & Cellular Proteomics, 4(12):2010–2021, 2005.

[3] Lily Ting, Ramin Rad, Steven P. Gygi, and Wilhelm Haas. MS3 eliminates ratio distortion in isobaric multiplexed quantitative proteomics. Nature Methods, 8(11):937–940, 2011.

[4] L. C. Gillet, P. Navarro, S. Tate, H. Rost, N. Selevsek, L. Reiter, R. Bonner, and R. Aebersold. Targeted Data Extraction of the MS/MS Spectra Generated by Data-independent Acquisition: A New Concept for Consistent and Accurate Proteome Analysis. Molecular & Cellular Proteomics, 11(6):O111.016717-O111.016717, 2012.

[5] Roland Bruderer, Oliver M. Bernhardt, Tejas Gandhi, Yue Xuan, Julia Sondermann, Manuela Schmidt, David Gomez-Varela, and Lukas Reiter. Optimization of Experimental Parameters in Data-Independent Mass Spectrometry Significantly Increases Depth and Reproducibility of Results. Molecular & Cellular Proteomics, 41(01):mcp.RA117.000314, 2017.

[6] Florian Meier, Philipp E. Geyer, Sebastian Virreira Winter, Juergen Cox, and Matthias Mann. BoxCar acquisition method enables single-shot proteomics at a depth of 10,000 proteins in 100 minutes. Nature Methods, page 1, 2018.

[7] Daniel A. Abaye, Frank S. Pullen, and Birthe V. Nielsen. Peptide polarity and the position of arginine as sources of selectivity during positive electrospray ionisation mass spectrometry. Rapid Communications in Mass Spectrometry, 25(23):3597–3608, 2011.

[8] Jürgen Cox, Marco Y. Hein, Christian A. Luber, Igor Paron, Nagarjuna Nagaraj, and Matthias Mann. Accurate Proteome-wide Label-free Quantification by Delayed Normalization and Maximal Peptide Ratio Extraction, Termed MaxLFQ. Molecular & Cellular Proteomics, 13(9):2513–2526, 2014.

[9] Andrew Thompson, Jürgen Schäfer, Karsten Kuhn, Stefan Kienle, Josef Schwarz, Günter Schmidt, Thomas Neumann, and Christian Hamon. Tandem mass tags: A novel quantification strategy for comparative analysis of complex protein mixtures by MS/MS. Analytical Chemistry, 75(8): 1895–1904, 2003.

[10] Stefka Tyanova, Tikira Temu, Pavel Sinitcyn, Arthur Carlson, Marco Y. Hein, Tamar Geiger, Matthias Mann, and Jürgen Cox. The Perseus computational platform for comprehensive analysis of (prote)omics data. Nature Methods, 13(9):731–740, 2016.

[11] Timothy Clough, Melissa Key, Ilka Ott, Susanne Ragg, Gunther Schadow, and Olga Vitek. Protein quantification in label-free LC-MS experiments. Journal of Proteome Research, 8(11):5275–5284, 2009.

[12] Yuliya Karpievitch, Jeff Stanley, Thomas Taverner, Jianhua Huang, Joshua N. Adkins, Charles Ansong, Fred Heffron, Thomas O. Metz, Wei Jun Qian, Hyunjin Yoon, Richard D. Smith, and Alan R. Dabney. A statistical framework for protein quantitation in bottom-up MS-based proteomics. Bioinformatics, 25(16):2028–2034, 2009.

[13] Ludger J.E. Goeminne, Andrea Argentini, Lennart Martens, and Lieven Clement. Summarization vs peptide-based models in label-free quantitative proteomics: Performance, pitfalls, and data analysis guidelines. Journal of Proteome Research, 14(6):2457–2465, 2015.

[14] Ludger J. E. Goeminne, Kris Gevaert, and Lieven Clement. Peptide-level Robust Ridge Regression Improves Estimation, Sensitivity, and Specificity in Data-dependent Quantitative Label-free Shotgun Proteomics. Molecular & Cellular Proteomics, 15(2):657–668, 2016.

[15] Hiromi W.L. Koh, Hannah L.F. Swa, Damian Fermin, Siok Ghee Ler, Jayantha Gunaratne, and Hyungwon Choi. EBprot: Statistical analysis of labeling-based quantitative proteomics data. Proteomics, 15(15):2580–2591, 2015.

[16] Robert Rosenthal. Combining results of independent studies. Psychological bulletin, 85(1):185, 1978.

[17] Jeremy D. O’Connell, Joao A. Paulo, Jonathon J. O’Brien, and Steven P. Gygi. Proteome-Wide Evaluation of Two Common Protein Quantification Methods. Journal of Proteome Research, 17 (5):1934–1942, 2018.

[18] Gordon K Smyth. Limma: linear models for microarray data. Bioinformatics and computational biology solutions using R and Bioconductor, pages 397–420, 2005.

[19] Simon Anders and Wolfgang Huber. Differential expression analysis for sequence count data. Genome Biology, 11(10):R106, 2010.

[20] Mark D. Robinson, Davis J. McCarthy, and Gordon K. Smyth. edgeR: A Bioconductor package for differential expression analysis of digital gene expression data. Bioinformatics, 26(1):139–140, 2009.

[21] Mark D. Robinson and Alicia Oshlack. A scaling normalization method for differential expression analysis of RNA-seq data. Genome Biology, 11(3):R25, 2010.

[22] Alexander Zien, Thomas Aigner, Ralf Zimmer, and Thomas Lengauer. Centralization: A new method for the normalization of gene expression data. Bioinformatics, 17:S323–S331, 2001.

[23] Jürgen Cox and Matthias Mann. MaxQuant enables high peptide identification rates, individualized p.p.b.-range mass accuracies and proteome-wide protein quantification. Nature Biotechnology, 26 (12):1367–1372, 2008.

[24] Jimmy K Eng, Ashley L McCormack, and John R Yates. An approach to correlate tandem mass spectral data of peptides with amino acid sequences in a protein database. Journal of the American Society for Mass Spectrometry, 5(11):976–989, 1994.

